# A Scalable, High-Throughput Optomotor Response Pipeline for Quantitative Analysis of Vision Using Infinity-Pools

**DOI:** 10.64898/2026.01.13.699264

**Authors:** Risa Suzuki, Damjan Kalšan, Norin Bhatti, Saul Pierotti, Gero Hofmann, Ewan Birney, Thomas Thumberger, Joachim Wittbrodt

## Abstract

The optomotor response (OMR) provides a robust readout assay for evaluating visually guided locomotion and has been widely applied to various animal models. We present a scalable, high-throughput OMR platform with tuneable and moving stripe patterns simultaneously projected to the side of 50 petri dishes. The setup is optimized for hatchling-stage medaka (*Oryzias latipes* and *Oryzias sakaizumii*) and larval stage zebrafish (*Danio rerio*) and integrates custom software for automated generation of visual stimuli and specimen detection, enabling precise and automated quantification of optomotor behaviour across thousands of individuals.

Using this platform, we systematically assess the visual performance of zebrafish and five medaka strains. By varying stripe width, direction, speed, contrast, and colour, we address spatial resolution, contrast, and colour sensitivity. Albino mutant medaka exhibited the highest sensitivity, responding at lower stripe widths and across a broader range of contrasts and chromatic conditions. Interestingly, some Cab and HO5 hatchlings displayed uni-directional response, revealing strain-specific visuomotor differences.

Our platform enables reliable detection of OMR behaviour already at hatching and supports robust analysis of visuomotor function. The modular design, automated detection, analysis pipeline, and capacity for large sample numbers make it a powerful tool for comparative vision research, high-throughput genetic screening, and systematic behavioural profiling.

## Introduction

Optomotor response (OMR) assays (Clark 1981) are non-invasive screening methods to assess visual and visuomotor function and to phenotype ocular or neurological disorders. OMR has been utilized in studying visual function (Brastrom et al., 2019; Mizoguchi et al., 2023; Neuhauss et al., 1999), social behaviour (Imada et al., 2010), and neural activities (Ahrens et al., 2013; Akiba et al., 2020) underlying motion detection and stabilization across different experimental models.

OMR is analysed by the locomotion pattern of an individual animal tested inside a defined arena, while a moving (stripe) pattern is displayed around or beneath it. Several variations of OMR setups are available that are tailored to the size and biology of the test animal in question. The most simple setup comprises a mechanically revolving slit-drum or a printed stripe pattern, the motion of which is recognized as a turning stripe pattern for the animal inside the test arena (Carvalho et al., 2002; Kretschmer et al., 2013; Matsuo et al., 2018). A drawback of these setups is the fixed pattern and their limited scalability for high-throughput.

More advanced setups employ computer or tablet displays projecting the moving pattern as video playback (Brastrom et al., 2019; Suzuki et al., 2024). This allows easy adjustment of the displayed pattern with respect to moving direction, velocity, contrast and colour. Parallel acquisition of multiple arenas was achieved by placing the display panel below the arena and filming from above (Brastrom et al., 2019; Neuhauss et al., 1999; Suzuki et al., 2024). However, this configuration bears problems to both, the biology of the test organism, as not all animals are robustly triggered by patterns displayed from below and the video analysis if the pattern is visible in the acquired movie (contrast challenges). In addition, uni-directed stripe patterns displayed from below or linear test arenas restrict the movement path of the test animal. Furthermore, the field of view of test animals differs. For example, a recent study in fish demonstrated that at a comparable larval stage, zebrafish (*Danio rerio*) primarily respond to motion from below, whereas medaka (*Oryzias latipes*) integrate motion over a broader visual field (Isoe et al. 2025). In addition, patterns displayed from below only project to the dorsal part of the retina, leaving the larger part without stimulus. Lateral projections in contrast cover most of the retinal tissue, likely engaging more neural circuits (Fig. 1A). These considerations underscore the need for adjustable OMR setups supporting high-throughput experimentation to enable comparative studies across species and genetic screens.

**Figure 1.**
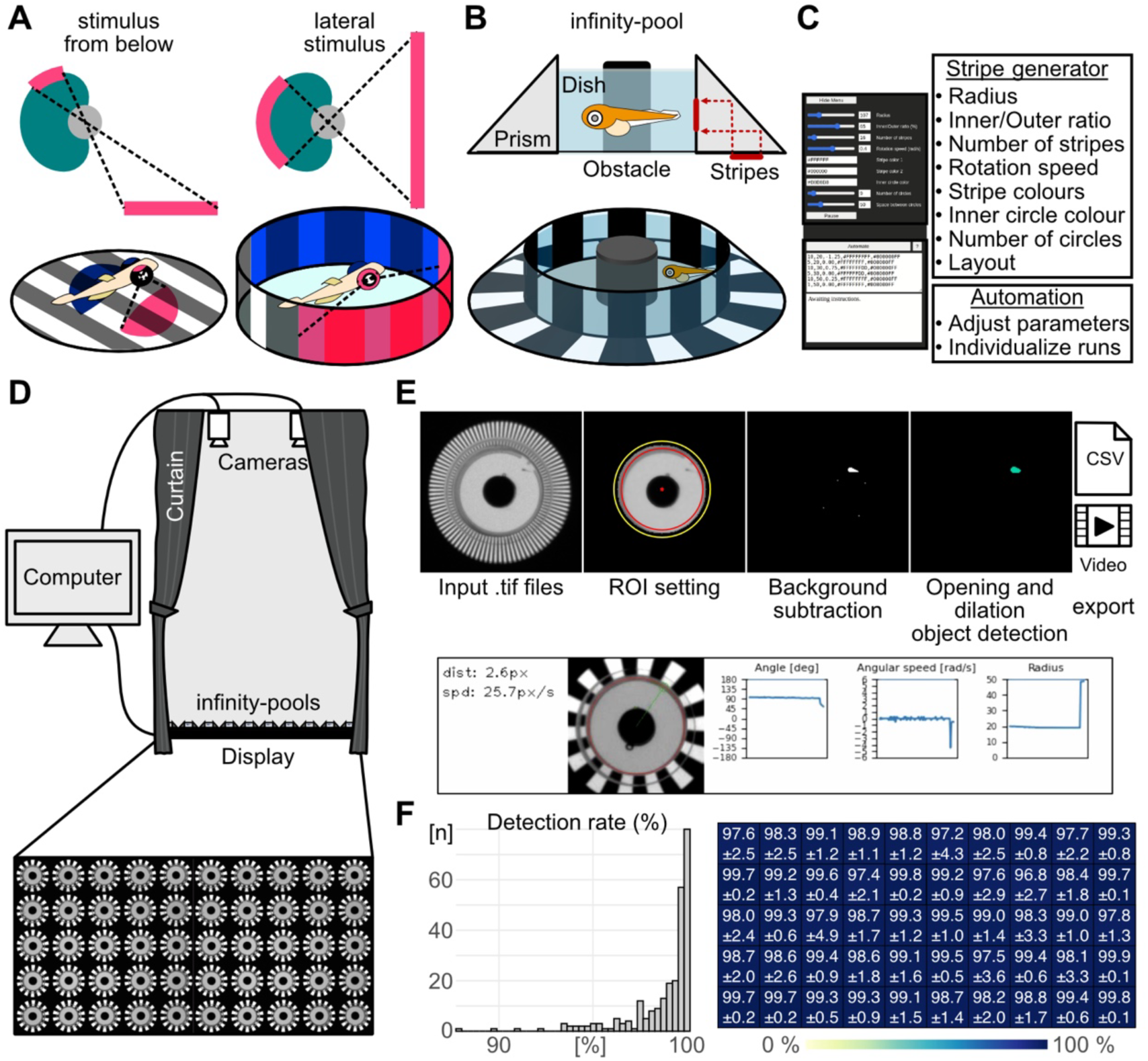
Optomotor response (OMR) and analysis pipeline of the infinity-pool setup. (A) Area of stimulus projected to retinal regions. Stimuli from below primarily project to the dorsal retina, whereas lateral stimuli engage both the dorso-ventral and naso-temporal axes. (B) The ‘infinity-pool’ comprises a 36 mm diameter Petri-dish with a surrounding plexiglass prism. The 45° slope projects patterns displayed below the arena laterally (red lines) to the hatchling. A black obstacle (12 × 10 mm) was positioned at the dish centre. (C) Stripe generator software provides parameter control to define the radial stripe pattern, layout and script-based animation. (D) High-throughput infinity-pool OMR setup: 50 infinity-pool units mounted on an upward-facing monitor displaying the stripe pattern, with cameras recording from above. To eliminate external visual stimuli, the entire setup is covered by black curtains. (E) Pipeline of the image processing workflow including manual or automatic selection of the region of interest (ROI), background subtraction, image processing, including opening and dilation, object detection (i.e. hatchling) followed by calculation of hatchling movement and exporting of data (CSV file) with optional video (Supplementary Movie 1) rendering. Example frame of an output video. Hatchling, green circle; ROI, red circle. The polar coordinates including angle (degrees) and radius (pixels, distance from the ROI centre), angular speed (radians/second), distance travelled between consecutive frames (pixels), and speed (pixels/second) are overlaid in real time. (F) Histogram of detection rate per dish calculated as the proportion of frames with hatchling-like objects, relative to total number of frames, based on 9 experiments with 1 day post hatch medaka. Mean detection rate per dish position. Data are presented as mean ± standard deviation (SD).

Here we describe an OMR setup designed to project the patterns to the arena wall that is simple to construct from standard components, readily adjustable in scale, and optimized for high-throughput acquisition and analysis. It enables fully parallelized assays (here up to 50) with precise control of visual parameters - including stripe width, contrast, colour, speed, and direction - each of which can be scripted for automated operation with minimal user intervention. The cylindrical arena employed, further offers an “uninterrupted” movement path of the test animal, allowing unlimited duration and flexible stimulus presentation, which increases the precision and reliability of phenotype quantification.

We used this setup to investigate differences in visual function, i.e. spatial, contrast and colour sensitivity of “classic” medaka inbred strains (Cab, HO5, HdrR; *Oryzias latipes*), Kaga (*Oryzias sakaizumii*), QuiH (pigment free albino mutants; *Oryzias latipes*) as well as zebrafish (*Danio rerio*). Using our setup, we found that QuiH hatchlings at 1 day post hatch (dph) reacted and adjusted their swimming speed and direction according to the projected revolving stripe pattern already at a thin stripe width. Cab hatchlings, in contrast, only did so at a wider stripe width. Five days post fertilization (dpf) zebrafish larvae responded to OMR stimuli at intermediate stripe widths. Moreover, QuiH hatchlings showed highest contrast and colour sensitivity. In addition, the overall response rate was higher in HdrR and QuiH whereas Cab and HO5 prominently exhibited uni-directional swimming behaviour, reflecting their visual capacity as well as behavioural differences.

## Results and Discussion

Small teleost fish, such as the Japanese medaka (Asai et al., 2011; Centanin & Wittbrodt, 2014) and zebrafish (Chhetri et al., 2014; Richardson et al., 2017), serve as excellent vertebrate models to study visuomotor function due to their short embryonic development, genetic tractability, small body size, and ease of maintenance (Iwamatsu, 2004; Kirchmaier et al., 2015; Wittbrodt et al., 2002). Availability of various genome editing tools and high fecundity enables cost-effective large-scale experiments for disease modelling (Gücüm et al., 2021; Ichimura et al., 2013), population genomic research (Gierten et al., 2025; Welz et al., 2025), and toxicological investigations (Dasmahapatra et al., 2023; Padilla et al., 2009).

We established an ‘infinity-pool’ setup for adjustable and high-throughput OMR. We achieve the lateral projection by a circular prism (Ring with 45° outer cut-off, Ø 56 mm outer diameter and Ø 36 mm inner diameter) made from acrylic glass (polymethyl methacrylate, PMMA; Fig. 1B) placed on a flat mounted 4K monitor. The 45° slope of the prism allows total internal reflection of the light emitted from the monitor below, thus projecting the stripe pattern displayed in a ring on the monitor onto the cylindrical wall of the Petri-dish placed in the centre of the prism (Fig. 1B). A black obstacle made of sandblasted polyoxymethylene (diameter/height = 12/10 (mm)) is placed at the centre of the dish to ensure that only one eye is exposed to the stimulus.

Projecting the stimulus onto the arena wall also facilitates easy detection of the specimens. The brightness in the swimming arena (central circle covered by the Petri dish) is uniformly adjusted to maximize the contrast to the specimen for efficient detection and tracking (Fig. 1; Supplementary Movie S1). To control the animated stimuli and background, we developed an open-source software tool (Stripe Generator; https://github.com/dkalsan/stripe-generator) that allows the adjustment of brightness, contrast, colour, stripe width, stripe number, as well as rotation speed and direction (Fig. 1C). All parameters, including the duration of the animation and pause intervals, can be scripted for full automation, eliminating the need for user intervention during experiments (Fig. 1C). The software also allows the adjustment of the layout and size of the arenas, i.e. the number of specimens that can be simultaneously tested on a single 4K monitor (Fig. 1D). In our case, one test arena is placed in a square with 6 cm side length. Using a 32-inch monitor, the (animated) stripe patterns can be multiplexed on the computer monitor, which allows for parallel projection and acquisition of 50 individual hatchlings with two cameras (CellCam Centro 200MR) mounted above the screen (1.8 m distance, Fig. 1D). In contrast to the fixed design of linear-pool-style-OMR assays (Suzuki et al., 2024) or traditional drum-style-OMR setups (Matsuo et al., 2018), our ‘infinity-pool’ OMR setup is fully flexible in terms of size of the swimming area, the visual cues and the number of arenas that can be adjusted to the needs of the assay and model system employed.

The ‘Fish Detector’ (https://github.com/dkalsan/fish-detector) - our custom-developed software - automatically classifies objects exceeding a predefined size threshold as specimens (Fig. 1E). For each video frame, the software extracts XY centroid coordinates and calculates angular movements of detected objects within the ROIs, thus enabling detailed tracking of hatchling behaviour. Output data are provided as CSV files and as annotated videos with overlaid tracking information (Fig. 1E; Supplemental Movie 1). When tested with 259 medaka hatchlings (1 dph), the detection algorithm reached high per-frame detection rates, typically 98-99%, with the lowest observed value being 96.9%. Detection rates were calculated as the proportion of frames in which hatchling-like objects were successfully identified independent of dish position (Fig. 1F).

To evaluate the robustness of the OMR setup, 20 hatchlings from the medaka Cab strain were tested in contrast to a negative control group of nine blinded hatchlings. Hatchlings were put into individual arenas and acquisition was initiated by a 5-minute acclimation period (Fig. 2A). Following this period, hatchlings were sequentially exposed to stripe stimuli varying in width, motion direction, and speed. Each of the 21 sets consisted of: a fixed stripe width and 2.5 minutes clockwise (CW) motion, a 30 seconds pause, 2.5 minutes of counterclockwise (CCW) motion, and a subsequent 30 seconds pause. For the subsequent set, the stripe width was increased (Fig. 2A).

**Figure 2.**
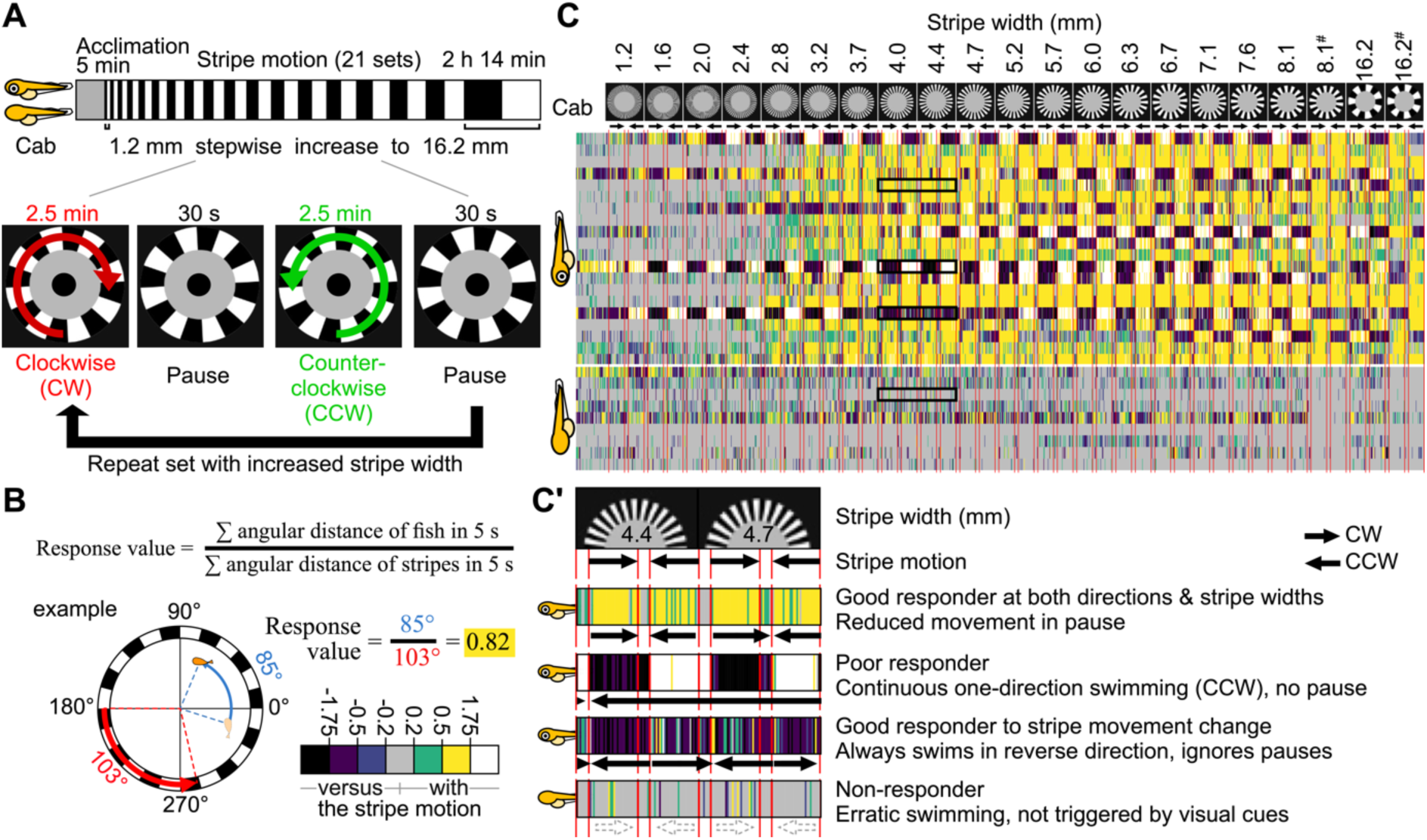
The infinity-pool setup robustly triggers OMR in medaka Cab strain. 1 day post hatch Cab medaka, with intact or removed eyes were used for optomotor response (OMR) evaluation. (A) Timeline of OMR experiment: after 5-minute acclimation, hatchlings were exposed to moving stripes (21 sets). Each set consisted of a fixed stripe width, 2.5-minute clockwise (CW) motion, 30-second pause, 2.5-minute counterclockwise (CCW) motion, and 30-second pause. Subsequent sets used progressively thicker stripes, ranging from 1.2 to 16.2 mm. (B) Response values were calculated every 5 seconds as the ratio of “angular distance swam by the hatchling” to “angular distance of stripe motion.” During pauses, the stripe speed from the preceding motion phase was used for calculations. Colour coding reflects response values indicating swimming against the stripe motion (black, purple; ≤ -0.5) via non-responding (blue, gray and green; from -0.5 to +0.5) to swimming with stripe motion (yellow, white; ≥ +0.5). (C) Heatmap of response values (columns) for each hatchling (rows) across stripe motions (direction indicated by arrows). Red lines mark onset/offset of stripe motion, i.e. pause. Stripe colours were black and white. Stripe speed was 20.6°/s, except where indicated with # (61.8°/s). (C’) Representative examples of hatchling swimming behaviour from C (black boxes).

To quantify the behaviour of hatchlings to the visual stimuli, a response value was calculated for 5 second bins as the ratio of the summarized angular distance moved by the hatchling divided by the movement of the revolving stripe during the same interval (Fig. 2B). Each 2.5-minute motion phase (clockwise [CW] or counterclockwise [CCW]) thus yields 30 response values per hatchling. A hatchling was considered responsive to a given stripe width, if in both the CW and CCW phases, at least 16 out of the 30 response values met or exceeded a predefined threshold. This threshold was set at response values ≥ 0.5 (indicating swimming with direction of stripe motion; green, yellow, white (Fig. 2C-C’)) or ≤ -0.5 (indicating swimming versus the direction of stripe motion; blue, purple, black (Fig. 2C-C’)). Moreover, directional consistency was required: hatchlings that followed the stripe motion in the CW phase had to do the same in the CCW phase, and those that swam against the motion in one phase were expected to continue doing so in the subsequent phase. This criterion ensured that the behaviour was induced by the visual stimuli and not by random swimming in a certain direction.

This stripe pattern motion was repeated with progressively thicker stripe widths per set and adjusted speeds (Fig. 2C). The stripe widths used (in mm) were: 1.2, 1.6, 2.0, 2.4, 2.8, 3.2, 3.7, 4.0, 4.4, 4.7, 5.2, 5.7, 6.0, 6.3, 6.7, 7.1, 7.6, 8.1, and 16.2 (Fig. 2B). All stripe patterns moved at a speed of 20.6°/s. For the stripe widths of 8.1 mm and 16.2 mm the second round of exposure was repeated with stripe widths revolving at triple speed (61.8°/s; Fig. 2C).

When tested with 20 individuals of the medaka Cab strain at 1 dph, the majority of the hatchlings started following the stripe motion from the width of 2.8 - 4.4 mm (n =16; yellow or white blocks; representative example in Fig. 2C’). Notably, a subset of individuals displayed continuous uni-directional swimming, irrespective of the directionality of the stripe movement (n = 2; purple or black and yellow or white alternating blocks; representative example in Fig. 2C’). Two individuals exhibited movement in the reverse direction to the stripe motion (purple and black blocks; representative example in Fig. 2C’). By contrast, swimming behaviour of blinded hatchlings was constant but erratic, clearly not triggered by the visual cues displayed (n = 8; gray blocks; representative example in Fig. 2C’).

Teleost species have individually adapted to their niches, have diverged from foraging to behavioural traits, and inhabit different parts of the waterbody (limnic, benthic) all of which may be reflected by the physiology of their visual system. To address the differences, we used the infinity-pool OMR setup comparing the visual sensitivity and swimming behaviour across teleost strains and species. We compared hatchlings (1 dph) of five medaka strains (*Oryzias latipes*: Cab, HO5, HdrR, and QuiH - albino mutant, and *Oryzias sakaizumii*: Kaga), to 5 dpf zebrafish larvae using the same parameters as before (Fig. 3A).

**Figure 3.**
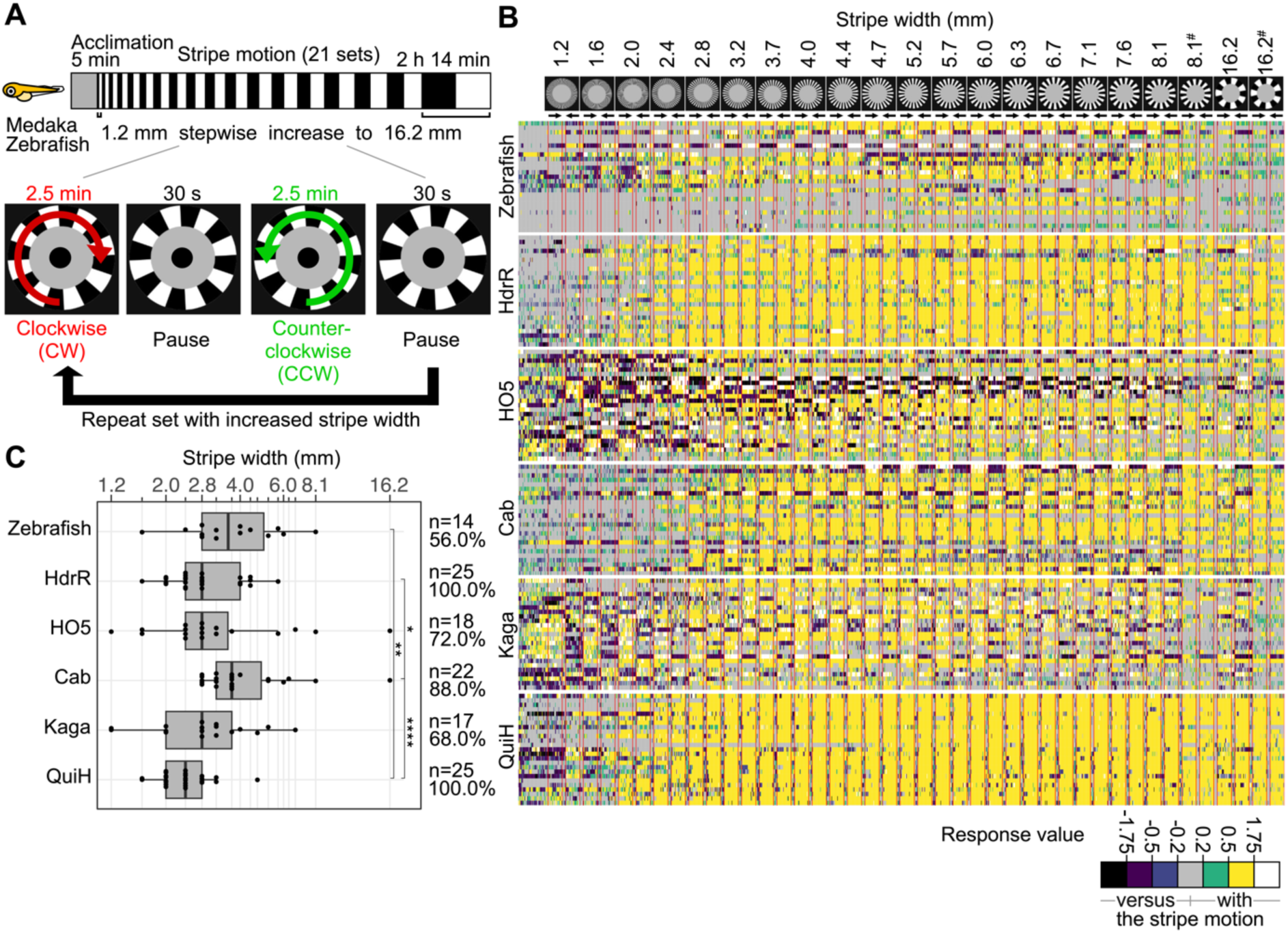
Behavioural and visual response parameters across five medaka strains and zebrafish in OMR experiments. (A) 1 day post hatch five medaka strains and five days post fertilization zebrafish were tested in the optomotor response (OMR) assay. After 5-minute acclimation, hatchlings were exposed to moving stripes. Each set consisted of a fixed stripe width, 2.5-minute clockwise (CW) motion, 30-second pause, 2.5-minute counterclockwise (CCW) motion, and 30-second pause. Subsequent sets used progressively thicker stripes, ranging from 1.2 to 16.2 mm. (B) Heatmap of response values (columns) for each hatchling (rows) across stripe motions (direction indicated by arrows). Colour coding reflects response values indicating swimming against the stripe motion (black, purple; ≤ -0.5) via non-responding (blue, gray and green; from -0.5 to +0.5) to swimming with stripe motion (yellow, white; ≥ +0.5). Red lines mark onset/offset of stripe motion, i.e. pause. Stripe speed was 20.6°/s, except where indicated with # (61.8°/s). (C) Box plots of the minimum stripe width that triggered response in individual hatchlings (dots) of each strain. The number (n) of responder hatchlings and their proportion (%) to the respective group are indicated. Statistical analyses were performed using Dunn’s tests with Bonferroni correction. Asterisks indicate level of statistical significance: * = p ≤ 0.05, ** = p ≤ 0.01, p = **** ≤ 0.0001.

The heatmap of the OMR analysis (Fig. 3B) revealed pronounced strain- and species-specific differences in visual performance at the individual level. Nearly all hatchlings of the HdrR and QuiH medaka strains robustly followed the stripe motions already at thin stripes, while the individual response in Cab, Kaga and HO5 as well as in zebrafish was more heterogeneous and the overall onset of response was shifted towards thicker stripes. Interestingly, Cab and HO5 hatchlings showed a higher prevalence of uni-directional swimming, characterized by sustained motion in a single direction (clockwise or counterclockwise) throughout both motion phases (Fig. 3B). Such behaviour may indicate reduced visual sensitivity, sensorimotor deficits, or diminished attentional control.

Analysis of the minimum stripe width eliciting OMR and the proportion of responsive individuals confirmed the strain- and species-specific differences in the onset of response (Fig. 3C). Among all groups, QuiH hatchlings exhibited the highest visual sensitivity, responding to the thinnest stripe width (median 2.4 mm). HdrR, HO5, and Kaga strains displayed comparable thresholds (median 2.8 mm), whereas Cab hatchlings required slightly thicker stripes (median 3.7 mm). Zebrafish required a median threshold of 3.6 mm. Responsiveness, measured as the fraction of individuals exhibiting any OMR, was high and robust across all medaka strains (≥87% in Cab, HdrR, and QuiH; 72% in HO5; 68% in Kaga). By contrast, 56% of zebrafish larvae displayed measurable OMR, reflecting developmental heterogeneity in visual system maturation and swimming competence at 5 dpf.

Having identified the optimal stripe width required to elicit a robust OMR, we next assessed contrast and colour sensitivity, factors likely more relevant to foraging and predator avoidance in natural environments. Following a 5-minute acclimation period, hatchlings were presented with sets of stripe motion at a fixed speed (20.6°/s) and stripe width (8.1 mm) under varying contrast and colour conditions. Stimuli included seven levels of grayscale contrast, colour contrast conditions pairing black with increasing intensities of blue, red, or green. In addition, we exposed the specimens to eight colour pairs previously identified as challenging to distinguish in human colour-blindness tests (orange/pink, pink/gray, pink/light blue, yellow/light green, light green/orange, gray/yellow, light blue/light green, and blue/purple; Fig. 4).

**Figure 4.**
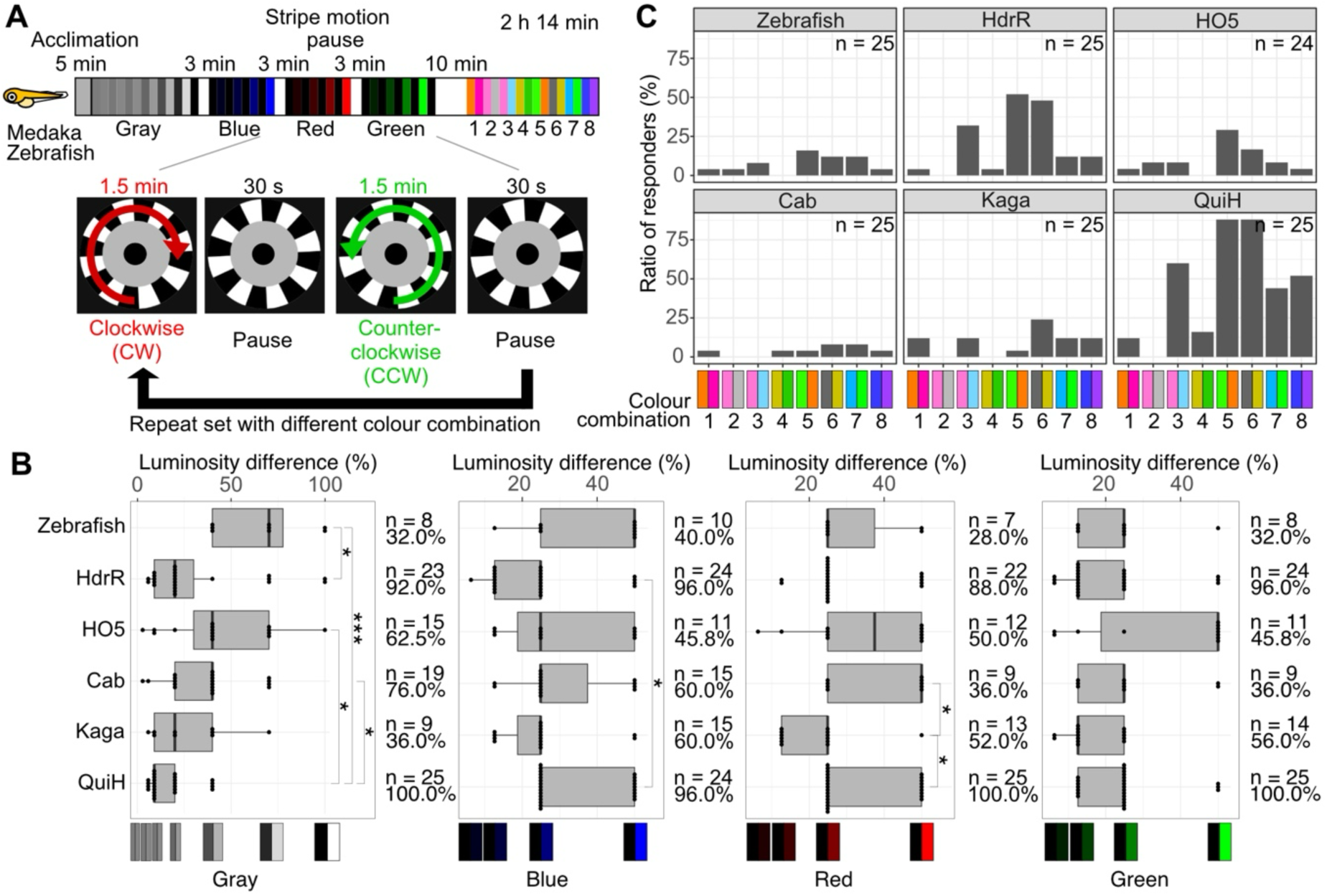
Contrast and colour sensitivity evaluation across medaka strains and zebrafish. (A) 1 day post hatch medaka from five strains and 5 days post fertilization zebrafish were tested in OMR assays with different stripe contrasts and colour conditions. After 5 minutes of acclimation, hatchlings were exposed to the stripe motions. Each set was performed with a fixed speed (20.6°/s) and stripe width (8.1 mm) and consisted of 1.5-minute clockwise (CW) motion, 30-second pause, 1.5-minute counterclockwise (CCW) motion, and 30-second pause. In subsequent sets the contrast or altered stripe colour combinations were as follows: gray/gray (7 combinational change from gray/gray to black/white stripes with luminosity differences of: 2%, 6%, 10%, 20%, 40%, 70%, and 100%), 3-minute pause; black/blue (4 differential blue luminosity: 6.3%, 12.6%, 24.9 %, 50%), 3-minute pause; black/red (4 differential red luminosity: 6.3%, 12.6%, 24.9 %, 50%), 3-minute pause; black/green (4 differential green luminosity: 6.3%, 12.6%, 24.9 %, 50%), 10-minute pause; 8 colour pairs previously identified as challenging to distinguish in human colour-blindness tests (combinations of stripes with the same luminosity; 1: orange/pink, 2: pink/gray, 3: pink/light blue, 4: yellow/light green, 5: light green/orange, 6: gray/yellow, 7: light blue/light green, 8: blue/purple). (B) Box plots of the lowest contrast that triggered response in individual hatchlings (dots) of each strain. The number (n) of responder hatchlings and their proportion (%) to the respective group are indicated. Statistical analyses were performed using Dunn’s tests with Bonferroni correction. Asterisks indicate level of statistical significance: * = p ≤ 0.05, ** = p ≤ 0.01, p = *** ≤ 0.001. (C) Histogram showing the proportion of responders to each challenging colour pair (1-8) identified from human colour-blindness tests per strain.

Each set consisted of 1.5 minutes of CW stripe motion, a 30 seconds pause, 1.5 minutes of CCW motion, and a final 30 seconds pause. Grayscale and primary colour tests were interleaved with 3 minutes rest periods, and a longer 10 minutes pause was implemented before human colour-blindness conditions (Fig. 4A). A hatchling was scored as responsive if > 9 out of 18 response values (absolute value) per phase reached or exceeded a threshold of 0.5 and the hatchling adapted to the changing motion directions.

When tested across seven levels of grayscale contrast (2%, 6%, 10%, 20%, 40%, 70%, and 100%), Cab, and HO5 hatchlings, as well as zebrafish larvae, displayed similar contrast sensitivity thresholds, with median values ranging from 40 - 70% contrast differences (Fig. 4B). HdrR and Kaga exhibited moderate sensitivity, with median threshold at 20% contrast difference (Fig. 4B). Conversely, QuiH hatchlings showed markedly enhanced contrast sensitivity, responding reliably at differences as low as 10% (median threshold) (Fig. 4B). This response distribution of the different strains and species largely follows their response to stripe width changes (Fig. 3).

In the black/colour combinations, the black/green set was most robustly responded to even at low luminosity contrast (Fig. 4B) across all species and strains except for HO5. In the black/blue combination, HdrR and Kaga strains exhibited a consistent high sensitivity and response to low luminosity differences while the other strains and species performed at a comparable level (Fig. 4B). Strikingly, the Kaga strain outperformed all other strains and species in response to black/red colour combinations (Fig. 4B). Considering the sensitive and robust response to low contrast black/green triggers in all species and strains, likely reflects the adaptation to a green natural environment.

In the human colour blindness test set, predominantly combinations including the green spectra (colour set 5, 6, 7) triggered the highest responses in each species and strains (Fig. 4C). This only holds true if the second colour is not green (colour set 4; Fig. 4C). In the strain/species comparison, QuiH hatchlings consistently outperformed all others, responding to subtle chromatic differences in most colour combinations, indicating their high contrast and colour discrimination capabilities.

Taken together, these results demonstrate functional differences in visual performance likely reflecting underlying genetic variation, shaped by natural selection in distinct ecological contexts. For example, the Kaga strain originates from a region with one of the shortest annual daylight durations in Japan, averaging 1603 hours for the years 2020 to 2024 (Japan Meteorological Agency database). This is considerably shorter than the average of 2234 hours recorded in Nagoya, the region of origin for the HdrR strain and may favour enhanced detection of subtle contrast differences. Similarly, the absence of ocular pigmentation in QuiH albino mutants may represent an alternative explanation to sensitize the visual system to subtle contrast changes.

Taken together, the infinity-pool OMR assay is a robust and versatile fully integrated platform for visual stimulation, detection and analysis, enabling the assessment of visual function in medaka and zebrafish hatchlings. The setup facilitates instant and reliable evaluation of strain- and species-specific differences in visual acuity, contrast sensitivity, and colour discrimination, and can readily be extended to other behavioural parameters such as baseline activity and swimming dynamics. Its sensitivity allows detection of both, subtle and pronounced visual characteristics, while minimal acclimation times and the scalability of the setup make it suitable for high throughput behavioural phenotyping and subsequent genetic screening.

Several physical limitations remain inherent to the current implementation, including screen resolution, image refresh rate (Hertz), and the sensitivity and resolution of the camera chip. These technical parameters define the lower limits of detectable visual stimuli and may be improved with future iterations of the setup.

The infinity-pool OMR assay offers broad applicability beyond the present study. It can be readily adapted for large-scale genetic or pharmacological screens, including CRISPR-based approaches to dissect the genetic architecture of visual acuity using resources such as the MIKK panel (Fitzgerald et al., 2022; Leger et al., 2022). Likewise, drug screens targeting ocular and neural disorders could exploit this platform to identify modulators of photoreceptor physiology, retinotectal circuit integrity, or locomotor responses. Diseases such as acute photoreceptor loss, macular degeneration, or retinitis pigmentosa may particularly benefit from such scalable *in vivo* assays.

Finally, the platform is not limited to medaka and zebrafish but can be adjusted to a wide range of hatchlings or small (non-aquatic) animals that respond to visual cues and can be kept individually. In this way, the infinity-pool OMR pipeline establishes a broadly applicable and adaptable framework for the quantitative assessment of visually induced behaviour, providing a basis for comparative, genetic, and translational research across (aquatic) model systems.

## Materials and methods

### Animal husbandry and ethics

Medaka and zebrafish were maintained in closed stocks at the fish facility of Heidelberg University under the supervision of the local representative of the animal welfare agency. Fish husbandry (permit number 35–9185.64/BH Wittbrodt) was performed in accordance with local animal welfare standards (Tierschutzgesetz §11, Abs. 1, Nr. 1) and following European Union animal welfare guidelines (Bert et al., 2016).

The following strains used in this study - *Oryzias latipes*: Cab strain (Wittbrodt et al., 2002), QuiH strain (Kirchmaier et al., 2015), HdrR strain (Spivakov et al., 2014), and HO5 strain (Spivakov et al., 2014); *Oryzias Sakaizumii*: Kaga strain (Asai et al., 2011; Spivakov et al., 2014); *Danio rerio*: AB strain [ZIRC, ZFIN: ZBD-GENO-960809-7] - were bred and maintained as previously described (Koster et al., 1997). HO5 (strainID: IB188) and HdrR-II1 (strainID: IB178) were supplied by NBRP Medaka (https://shigen.nig.ac.jp/medaka/).

All experimental procedures were performed according to the guidelines of the German animal welfare law and approved by the local government (Tierschutzgesetz §11, Abs. 1, Nr. 1, husbandry permit number 35-9185.64/BH Wittbrodt).

Medaka hatchlings 1 day post hatch (stage 40) (Iwamatsu, 2004) and zebrafish larvae at 5 days post fertilization were both used before the initiation of self-feeding (Hernandez et al., 2018; Watanabe et al., 2023). They were maintained at 28°C on a 14 hours light/10 hours dark cycle in Embryo Rearing Medium (ERM) (17 mM sodium chloride, 0.4 mM potassium chloride, 0.27 mM calcium chloride dihydrate and 0.66 mM magnesium sulphate heptahydrate at pH 7) for medaka and E3 medium (5 mM NaCl, 0.17 mM KCl, 0.33 mM CaCl_2_, 0.33 mM MgSO_4_) for zebrafish.

### Preparation of blinded hatchlings

0 dph hatchlings (pre-self-feeding fry (Watanabe et al., 2023)) were anesthetized in 0.02% Tricaine for 5 minutes until swimming ceased, after which the eyes were surgically removed using forceps (DUMONT®, Cat: 0208-5-PO & 0208-55-PO, Inox 08). Following surgery, the hatchlings were monitored, and those swimming normally the next day were used for subsequent OMR.

### Infinity-pool optomotor response (OMR) setup and protocol

The OMR setup is illustrated in Figure 1. The frame of the setup was constructed using extruded aluminium profiles (30 x 30 mm), assembled with electro-galvanized and passivated steel nuts and bolts. A black curtain was installed around the frame to minimize external light interference. The display (Gigabyte M28U 71,1 cm (28”) 4K Ultra HD LED | Huawei) was disassembled to allow spatial separation of the screen and its electronic components, thus reducing the heat transfer to the display surface. Each test arena consisted of a custom-made prism (outer/inner diameter = 56 mm/36 mm) made of polymethyl methacrylate and a 36 mm polystyrene dish (SARSTEDT 82.1135.500) with a black obstacle (diameter/height = 12/10 (mm)) made of sandblasted polyoxymethylene centrally attached using LOCTITE® 460 adhesive and a custom mold (Supplementary Fig. S2). 50 dishes with prisms were placed on the disassembled monitor.

Two CellCam Centro 200MR cameras, equipped with Goyo Optical lenses (GM12HR41216MCN), were mounted 1.8 meters above the setup to record behaviour. Video acquisition was performed using μManager software (Micro- Manager-2.0.0-gamma1-20210214). Visual stimuli were generated using the custom developed Stripe generator software (https://github.com/dkalsan/stripe-generator). Upon hatching, specimens were transferred to fresh Petri dishes with 1x ERM in 28 °C incubator until the following day for the OMR testing at 1 dph for medaka or 5 dpf for zebrafish.

On the day of the experiment, all hatchlings were screened under a binocular microscope and selected based on their swimming ability and wild-type morphology. OMR experiments were performed during the day time (9:00-17:00). All behavioural tests were conducted at room temperature (21 - 23 °C).

### Stripe Generator configuration

Stripe motion was generated using Stripe Generator developed using p5.js (version 1.4.1). Further details are available at (https://github.com/dkalsan/stripe-generator). The stripe configuration used for parallelizing 50 arenas on the display was as shown in Table 1.

**Table 1.** Parameters used on Stripe Generator.

| Value | Parameter |
| --- | --- |
| 92 | Radius |
| 64 | Inner/Outer ratio (%) |
| 7 | Number of stripes |
| 0 | Rotation speed (rad/s) |
| #FFFFFF | Stripe colour 1 |
| #000000 | Stripe colour 2 |
| #B8B8B8 | Inner circle colour |
| 50 | Number of circles |
| 5 | Space between circles |

OMR test with varying stripe width: each trial began with a 5-minute acclimation phase: 1 minute with a black background followed by 4 minutes with a stationary version of the first stripe pattern. Hatchlings were then exposed to moving stripe stimuli, with varying motion direction, stripe width, and speed. Each stimulus set included 2.5-minute clockwise (CW) motion, a 30-second pause, 2.5-minute counterclockwise (CCW) motion, and another 30-second pause. This sequence was repeated with progressively thicker stripe widths: 1.2, 1.6, 2.0, 2.4, 2.8, 3.2, 3.7, 4.0, 4.4, 4.7, 5.2, 5.7, 6.0, 6.3, 6.7, 7.1, 7.6, 8.1, 8.1, 16.2, and 16.2 mm (Fig. 2A). Stripe motion speed was set to 20.6°/s, except for the second round of the 8.1 mm and 16.2 mm conditions, which were presented at 61.8°/s. OMR test with varying colour and contrast: following a 5-minute acclimation period, hatchlings were subjected to a series of stripe motion stimuli at a fixed width (8.1 mm) and speed (20.6°/s). Each stripe motion set consisted of 1.5-minute of CW motion, a 30-second pause, 1.5-minute of CCW motion, and a second 30-second pause. The visual stimuli were presented in the following order, grayscale contrast series: 7 combinational changes from gray/gray to black/white stripes with luminosity differences of 2%, 6%, 10%, 20%, 40%, 70%, and 100%, by incrementally darkening one stripe and brightening the other. Colour contrast series: black paired with blue, red, or green with 4 different luminosities for each colour of 6.3%, 12.6%, 24.9 %, 50%, with a 3-minute pause between each colour set. Colour-pairs: after a 10-minute pause, hatchlings were exposed to eight colour combinations with the same luminosity known to be difficult to differentiate in human colour vision deficiencies: orange/pink, pink/gray, pink/light blue, yellow/light green, light green/orange, gray/yellow, light blue/light green, and blue/purple. 5

Throughout this study, “stripe width” refers to the combined width of one black and one white stripe.

### Tracking and analysis

Movement of hatchlings was tracked using a custom detection software “Fish detector”, developed in Python 3.8 and incorporating the following major packages: OpenCV, SciPy, NumPy, pandas, and Matplotlib. Detailed description of the software architecture and functionality is provided in (https://github.com/dkalsan/fish-detector). TIFF image stacks from the two-camera setup were separated using the above software into two stacks, each originating from one of the cameras. Regions of interest (ROIs) were assigned either automatically or manually. For manual ROI assignment, a Fiji macro was used to mark the centre of each circular arena and XY coordinates were recorded. These coordinates were used to generate a configuration file (config.json) for use with the Fish detector developed in this study.

In Fiji, the image analysis pipeline involved background subtraction followed by morphological operations (opening and dilation) to isolate objects. Objects exceeding a specified size threshold (≥5 pixels in area) were automatically classified as hatchlings. To reduce false detections, hatchling locations were further validated over video frames using a centroid tracking algorithm and simple post-processing techniques. Finally, for each frame, the software extracted both pixel and polar coordinates of the centroid of detected hatchling. Outputs were saved as CSV files, which were further processed using custom R scripts for behavioural phenotype quantification.

To extract timing of stripe motion transitions, mean gray value differences between adjacent frames were calculated using a Fiji macro. In short, the macro duplicates the frame sequence offset by one frame to create two stacks (‘Stack-1’ and ‘Stack-2’), and generates a differential stack using Fiji’s Image Calculator. The mean gray value per slice was measured, compiled, and saved as a CSV file.

Custom R scripts were used to identify stripe motion transitions, i.e. gray value differences below 0.8 for ≥25 consecutive frames were classified as stationary periods. Large gaps between these periods (Δ > 1500 frames) marked stripe motion onsets and offsets. A cumulative time column (Time) was generated using the calculated frame duration and frame count, enabling accurate alignment of behavioural data to stimulus timing.

Custom R scripts were used to calculate response values in 5-second bins. Hatchling movement was defined as the sum of angular changes; stripe motion was calculated as the product of stripe speed (°/s) and time (5 s). Positive response values indicated swimming in the same direction as stripe motion, negative values indicated the opposite. Stripe speeds were set at 20.6°/s (CW = positive; CCW = negative). For stationary periods (‘pauses’), either 20.6°/s was used or the speed of the previous phase, if different. Data with a detection rate below 75% were excluded from further analysis and plotting.

A hatchling was classified as responsive if it met the following criteria:

- For stripe width experiments: at least 16 response values (≥0.5 or ≤−0.5) in both CW and CCW phases.
- For contrast/colour experiments: at least 10 response values (≥0.5 or ≤−0.5) in both CW and CCW phases.

In addition, the hatchling had to show a change in swimming directionality between the CW and CCW phases. The minimum stripe width or luminosity differences eliciting a valid response was recorded per individual.

The response rate was defined as the proportion of hatchlings per group that responded to any stripe motion. Uni-directional swimming was defined as significant directional responses (≥20 counts of response values ≥0.5 or ≤-0.5) occurring in only one direction across both CW and CCW motion phases at the same stripe width. This was interpreted as a failure to respond appropriately to direction changes. Occurrence of one-directional swimming for each individual was counted and plotted. Detection rates were calculated as the proportion of non-missing frames relative to the expected frame count for each dish position.

### Data and code availability

All scripts used for the analysis are available on GitHub:

- Stripe generator: https://github.com/dkalsan/stripe-generator
- Fish detector: https://github.com/dkalsan/fish-detector
- Stripe Generator Automation Settings, Macros, Python scripts, and R notebooks used for automated detection, metadata handling, behavioural response analysis, and plotting: https://github.com/risasuzuki18/OMR-analysis (coding of these scripts was assisted by ChatGPT 4o and 5)

### Data visualization

Data visualization and figure assembly were performed using Fiji (Schindelin et al., 2012), ggplot2 (Wickham, 2016) in RStudio (Team R, 2020), Autodesk Inventor 2024 and Affinity Designer 2.6.3.

## Acknowledgement

We are deeply grateful to current and previous members of the Wittbrodt and Birney labs for their critical, insightful and constructive feedback on both the procedure and the manuscript. We especially thank S. Ansai for valuable discussions on improving medaka responsiveness to the OMR stimulus. We further thank N. Grammling for valuable support in assembling the setup and M. Majewski, E. Leist, R, Lipp, S. Erny and A. Saraceno for excellent fish husbandry.

