## Supplementary Material for "A Scalable, High-Throughput Optomotor Response Pipeline for Quantitative Analysis of Vision Using Infinity-Pools"

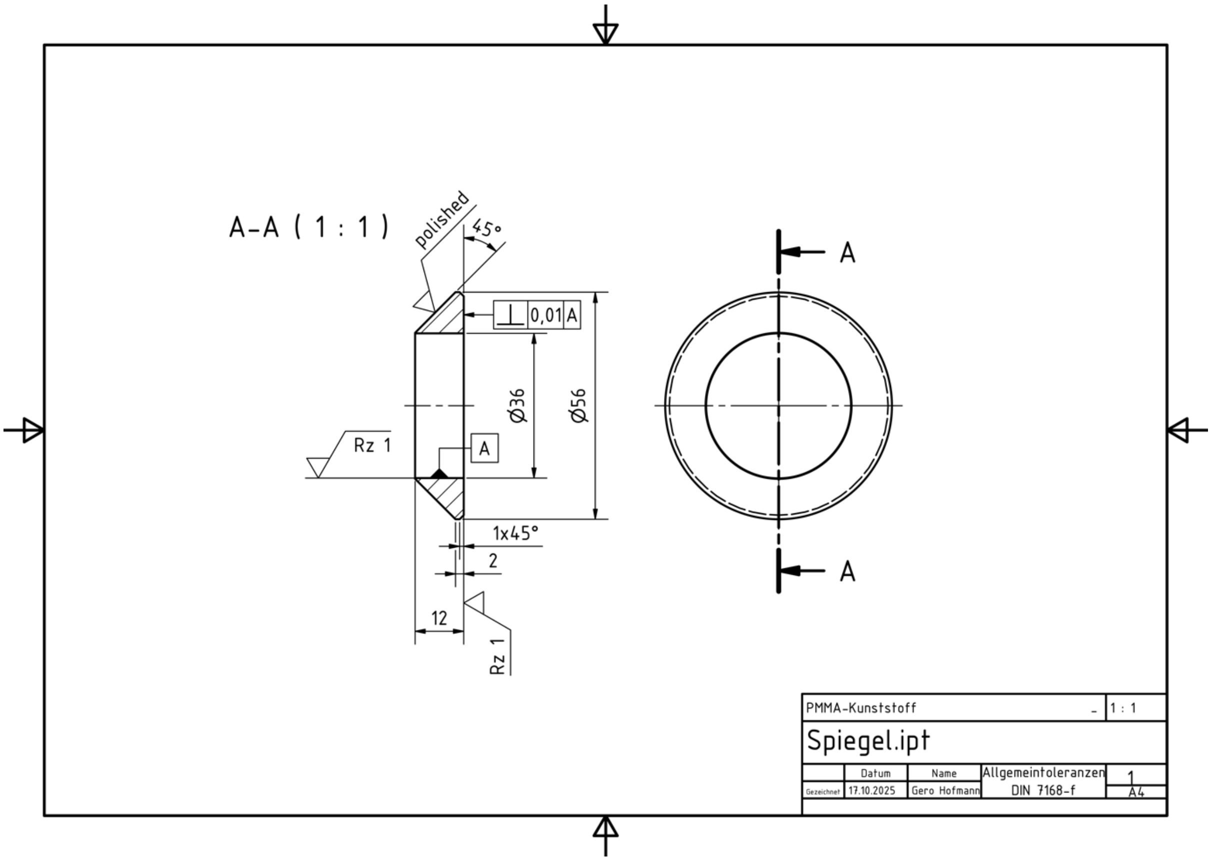


**Supplementary Figure 1 Technical drawing of the PMMA prism**

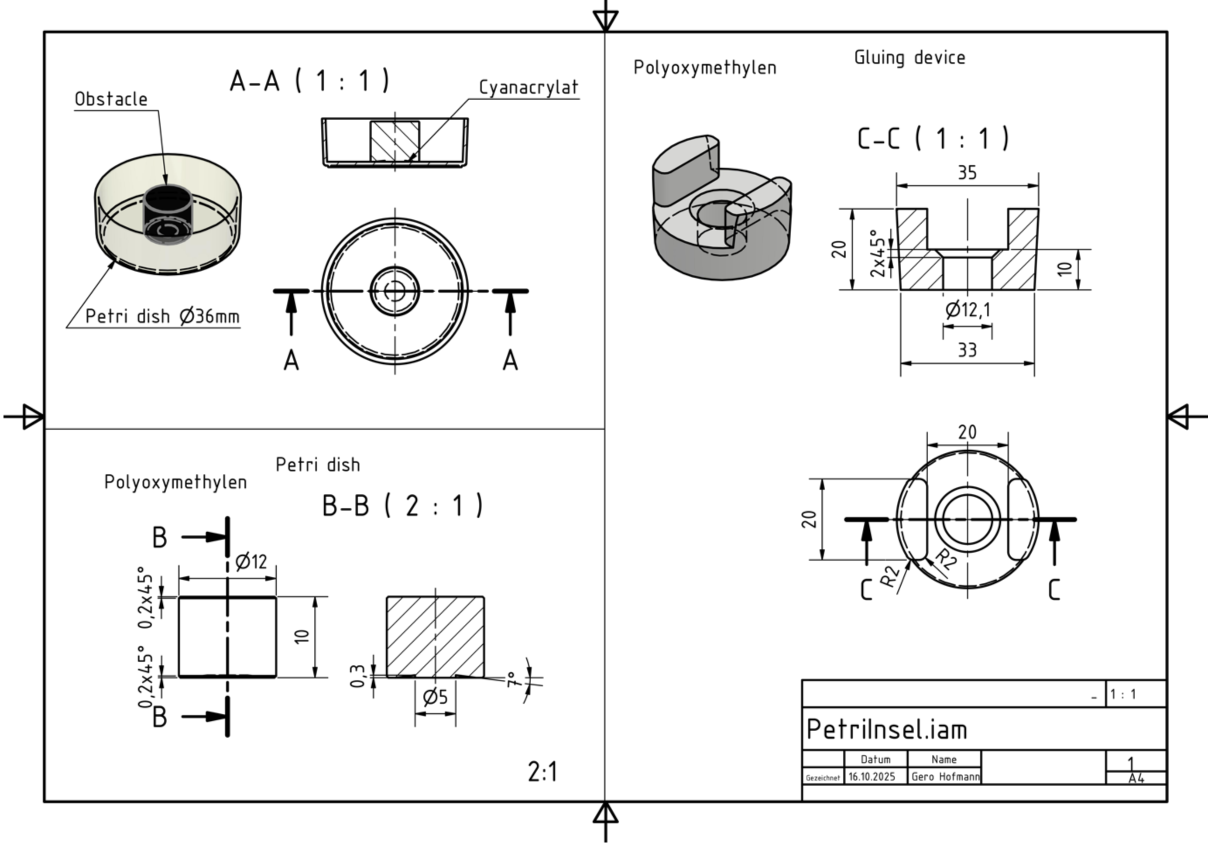


**Supplementary Figure 2 Technical drawing of the black obstacle and custom mold for its placement**

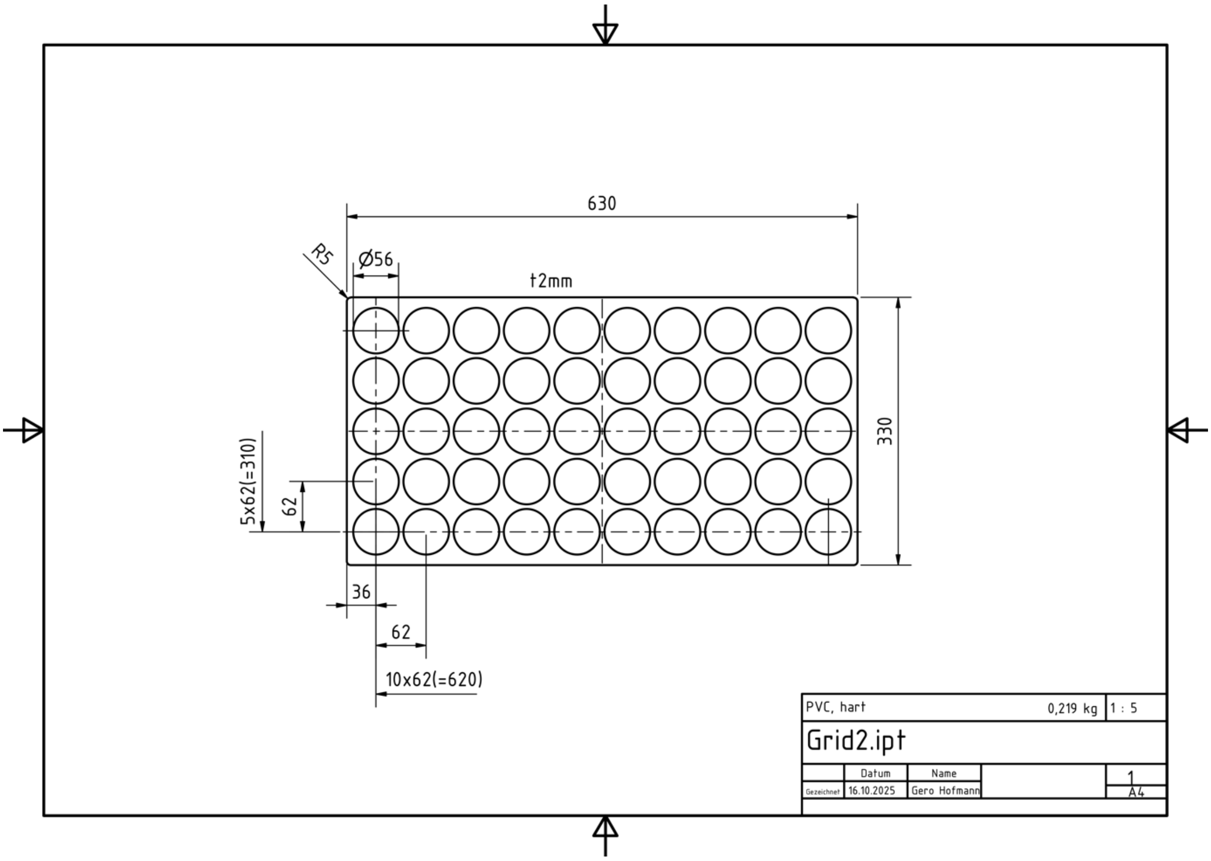


**Supplementary Figure 3 Technical drawing of the prism holder**

**Supplementary Movie 1 Example output from the Fish detector software showing responses of a single HdrR hatchling exposed to different stripe widths.**

Combined video clips of the behavior of a single 1 day post hatch HdrR hatchling in response to selected stripe widths within one experiment. Stripe width 1.2 mm: clockwise (CW) stimulus, no response. Stripe width 1.6 mm: counterclockwise (CCW) stimulus, onset of response. Stripe width 4.7 mm: example of pausing and resuming following the stripe motion. Stripe width 16.2 mm: 3× speed, continuous following of faster stripe motion. All clips were trimmed from the detection software output and vertically combined here.
